# Phosphoproteomics of ATR Signaling in Prophase I of Mouse Meiosis

**DOI:** 10.1101/2021.04.13.439649

**Authors:** Jennie R. Sims, Vitor M. Faça, Catalina Pereira, Gerardo A. Arroyo-Martinez, Raimundo Freire, Paula E. Cohen, Robert S. Weiss, Marcus B. Smolka

## Abstract

During mammalian meiosis, the ATR kinase plays crucial roles in the coordination of DNA repair, meiotic sex chromosome inactivation and checkpoint signaling. Despite the importance of ATR in meiosis, the meiotic ATR signaling network remains largely unknown. Here we defined ATR signaling during prophase I in mice. Quantitative analysis of phosphoproteomes obtained after genetic ablation of the ATR-activating 9-1-1 complex or chemical inhibition of ATR revealed over 12,000 phosphorylation sites, of which 863 phosphorylation sites were dependent on both 9-1-1 and ATR. ATR and 9-1-1-dependent signaling was enriched for S/T-Q and S/T-X-X-K motifs and included proteins involved in DNA damage signaling, DNA repair, and piRNA and mRNA metabolism. We find that ATR targets the RNA processing factors SETX and RANBP3 and regulate their localization to the sex body. Overall, our analysis establishes a comprehensive map of ATR signaling in spermatocytes and highlights potential meiotic-specific actions of ATR during prophase I.

## Introduction

Meiosis is a specialized cellular process whereby a single round of DNA replication is followed by two successive rounds of cell division to produce haploid gametes. To ensure proper chromosome segregation at the first meiotic division, meiotic cells must undergo a series of highly regulated processes, including programmed double-strand break (DSB) formation, recombination, and chromosome synapsis^1, 2^. The serine/threonine-protein kinase ATR (ataxia telangiectasia and Rad-3 related protein) has well-characterized roles in maintaining genome stability in mitotic cells^3–5^. In mammals, ATR also plays an essential role in spermatogenesis by promoting meiotic sex chromosome inactivation (MSCI), a process that is required for silencing of the X and Y chromosomes^6–8^. Impairment of ATR activity results in insufficient MSCI and germ cell elimination at the mid-pacyhytene stage of prophase I^7–13^. A major readout of ATR activity during MSCI is the phosphorylation of the histone variant H2AX within the dense heterochromatin domain of the nucleus that houses the X and Y chromosomes known as the sex body. Additionally, ATR regulates the sex body localization of several other DNA damage response proteins such as MDC1, CHK1 and BRCA1 in a feed-forward mechanism that allows these proteins to spread from the chromosome cores to the chromatin loops, ultimately resulting in MSCI^9, 14–19^. Furthermore, loss of ATR protein in spermatocytes results in defects in DSB repair and chromosome synapsis, implying that ATR regulates several aspects of meiotic progression^14, 20^. Despite the importance of ATR in meiosis, the mechanisms by which meiotic ATR signaling coordinates meiotic progression remains limited due to the complexity and interdependence of meiotic DNA repair, chromosome synapsis, and silencing of unsynapsed chromatin.

While ATR activation is less understood in meiotic cells than in somatic cells, it is likely that at least some of the molecular determinants of ATR activation are shared. In mitotic cells, ATR is activated at sites of single-stranded DNA (ssDNA) that arise during replication or DNA repair^3, 21^. ATR is recruited to RPA-coated ssDNA via interaction with ATRIP (ATR interacting protein)^22^, while the 9A-1-1 (RAD9A-RAD1-HUS1) checkpoint clamp is independently loaded at the dsDNA-ssDNA junction by the RAD17-RCF clamp loader^23, 24^. The ATR activating protein, TOPBP1 (topoisomerase binding protein 1), then interacts with the c-terminal tail of RAD9A and directly activates ATR via its ATR activation domain (AAD)^25–28^. Recently a second ATR activating protein, ETAA1 (Ewing’s tumor-associated antigen 1), was identified^29–31^. ETAA1 can directly activate ATR at RPA-coated ssDNA at replication forks and is thought to be important for ATR activation during unchallenged DNA replication^32, 33^. Once activated, ATR preferentially phosphorylates proteins at S/T-Q motifs^34^. In mitotic cells, ATR-mediated phosphorylation of H2AX and the MDC1 scaffold help promote checkpoint activation through ATR phosphorylation of the CHK1/CHK2 kinases, resulting in checkpoint-mediated cell cycle arrest^35, 36^. Meiotic roles for TOPBP1 in promoting key ATR signaling outcomes have been described^19^. Conditional TOPBP1 depletion in germ cells resulted in defective meiotic sex chromosome silencing as well as loss of ATR and H2AX phosphorylation in the sex body. The results suggest a prominent role for TOPBP1 in mediating the strong ATR signaling observed during pachynema. Intriguingly, this strong induction of ATR activity observed during normal spermatogenesis is compatible with the progression of the meiotic cell cycle. How ATR signaling in meiotic cells coordinates meiotic progression without imposing a checkpoint arrest remains a fundamental unanswered question.

While hundreds of targets of ATR have been characterized using quantitative phosphoproteomics mitotic cells^32, 37^, much less is understood about ATR signaling in meiosis. A comprehensive dataset of meiotic ATR-dependent phosphorylation events is necessary to further our understanding of the mechanisms by which ATR coordinates DNA repair, chromosome synapsis, checkpoint and MSCI pathways during meiosis. To define the network of phosphorylation events mediated by ATR in meiotic cells we performed extensive phosphoproteomic analyses of testes derived from mice with two independent methods of impairing ATR signaling. Given that ATR depletion results in embryonic lethality^35, 38, 39^, we used a genetic model of impairing ATR signaling whereby the 9-1-1 component RAD1 is conditionally depleted under the germ-cell specific *Stra8-Cre*. Depletion of RAD1 in germ cells is anticipated to disrupt all potential 9-1-1 complexes, including those that could form with the testes-specific paralogs *Hus1b*^40^ and *Rad9b*^41^, and therefore significantly impair ATR signaling^42^. In parallel, we collected testes from mice treated with ATR inhibitor (AZ20) and vehicle treated controls. We processed these tissues to isolate phosphorylated peptides, labeled them with amino reactive tandem mass tag reagents (TMT) followed by mass spectrometry to generate phosphoproteomic datasets. By combining these datasets, we were able to identify a set of over 800 high-confidence meiotic phosphorylation events that are dependent on both ATR and RAD1, including phosphorylation of crucial DNA damage signaling and DNA repair proteins. Intriguingly, we also detected ATR-dependent phosphorylation of several RNA metabolic proteins and found that that ATR-dependent phosphorylation controls the localization of key RNA processing factors. Overall, our analysis establishes a comprehensive map of ATR signaling in spermatocytes and highlights potential meiotic-specific actions of ATR during prophase I progression.

## Results

### A pharmacological and genetic approach to map ATR-dependent signaling in spermatocytes

Despite the central role of ATR in Prophase I of mammalian meiosis, the signaling network mediated by ATR remains largely unknown. Here, we generated phosphoproteomic databases from two sets of mice with independent methods of ATR inhibition. First, to genetically impair ATR activity, we utilized a *Rad1* conditional knockout model in which RAD1 is depleted in germ cells under a *Stra-8-Cre* promoter. Next, to chemically inhibit ATR, we treated mice with the ATR inhibitor AZ20. By comparing the databases generated from these two methods of ATR inhibition, we can take advantage of the tissue-specificity of the conditional knockout model as well as the acute inhibition of the kinase after ATR inhibitor treatment to identify a set of high-confidence ATR and RAD1-dependent phosphorylation events and overcome limitations arising from analysis of these two models individually (Fig. 1A).

**Figure 1.**
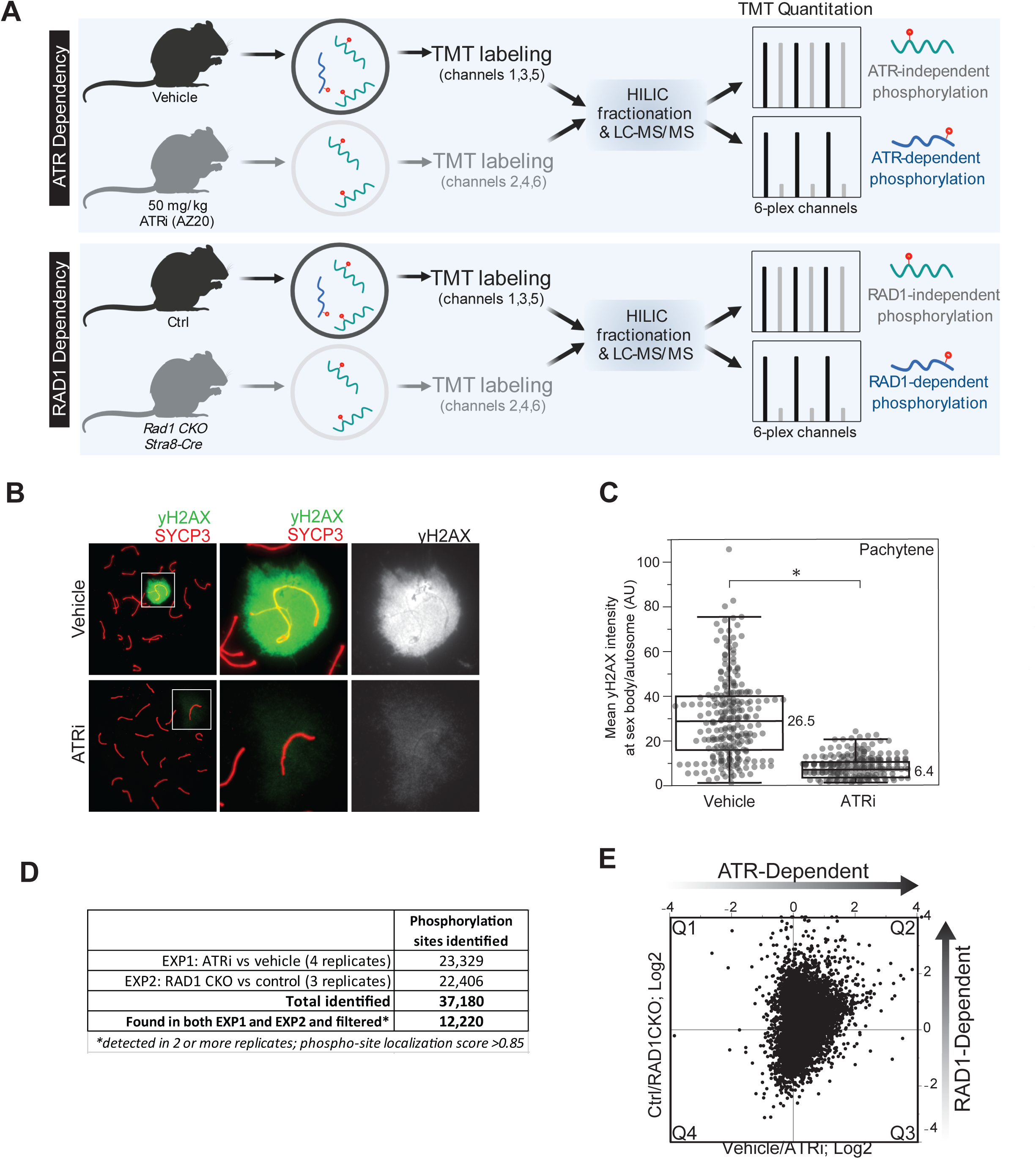
Approach for identifying ATR-dependent and RAD1-dependent phosphorylation events in mouse testes. A) Whole, decapsulated testes were collected from vehicle and AZ20-treated mice (top) or *Rad1 CKO* and control litter mates (bottom) and subjected to quantitative phosphoproteomic analysis to identify ATR-dependent and RAD1-dependent phosphorylation events (see methods for details). B) Immunofluorescence staining of meiotic spreads from ATRi or vehicle treated mice collected 4 hours after 50 mg/kg treatment with AZ20. C) Quantification of yH2AX intensity (4 vehicle mice, n=246 cells; 4 ATRi mice, n=309 cells p=0.019 measured by student’s t-test) (see methods for more details). D) Description of replicates and filters utilized in generation of the final phosphoproteomic dataset. E) Scatter plot of the final consolidated phosphoproteomic dataset.

We collected tissue from mice treated with ATR inhibitor AZ20 (ATRi), or vehicle, as well as from *Rad1* CKO and control mice (Fig. 1A). For ATRi, we treated mice with 50 mg/kg AZ20 per day and collected tissue at 2.5 or 3 days, following conditions previously described^7^. We also collected testis from mice 4 hours after one dose of 50mg/kg of AZ20. To ensure the ATRi dosage and timing was sufficient to impair ATR activity, we examined meiotic chromosome spreads to monitor the localization of yH2AX at the sex body during pachynema, which is dependent on ATR^2, 9, 43–47^. In mice treated with ATRi, we observed robust reduction in sex body yH2AX localization in pachynema staged cells 4 hours after a single dose of 50mg/kg treatment with AZ20 (Fig. 1B-C, S1A-B), indicating that ATR activity is rapidly impaired following AZ20 administration. We collected tissue for further mass spectrometry analysis from one pair of experimental and control mice under each of the treatment conditions for a total of four ATRi and vehicle control pairs of mice (2 pairs after 2.5-3 days of treatment and 2 pairs 4 hours after a single dose). We also collected decapsulated whole-testes from three *Rad1* CKO mice and litter-mate controls. Tissue samples were lysed and digested with trypsin, followed by enrichment of phosphopeptides and labeling with the 6-plex Tandem Mass Tag (TMT) reagent (Fig. 1A). HILIC pre-fractionation of TMT-labeled phosphopeptides allowed in-depth quantitative phosphoproteomic analysis, resulting in a total of 37,180 phosphorylation sites identified between the seven experiments (Fig. 1D). After selecting for high-quality phosphosites (localization score to > 0.85) and considering only phosphopeptides identified in both the AZ20 and *RAD1* CKO datasets in at least two independent samples, our final list yielded 12,220 quantitated phosphorylation sites (Fig. 1D-E, table S1).

### RAD1-dependent ATR signaling targets proteins involved in nucleic acid metabolism, DNA damage response and the cell cycle

As previously described, quantitative phosphoproteomic analysis combining datasets from chemical (ATRi) and genetic (*Rad1*-CKO) ATR inhibition was predicted to enrich for acute ATR signaling events specifically in germ cells. We therefore focused on phosphopeptides displaying consistent reduction in abundance in testes from both ATRi treated and *Rad1*-CKO mice (positioned in Q2, highlighted in Fig 2A). To address errors in quantitation and variation due to sample handling we applied the “Bow-tie” filter by excluding data points that fell outside of a fourfold of the log2 scale interval of correlation as previously described^48^. By comparing the number of phosphopeptides in each of the regions (quadrants) of the experimental correlation plots, we observed a biased distribution with approximately 4-fold more differentially phosphorylated peptides in Q2 compared to each of the other quadrants indicating that the primary mode of ATR activation during meiosis is RAD1 dependent (Fig. 2B, S4A). Gene enrichment analysis of each quadrant revealed that Q2 is enriched for gene ontology categories such as nucleic acid metabolism, DNA damage response and cell cycle (Fig. 2C, table S2), consistent with the expected roles for ATR and RAD1 in these processes.

**Figure 2.**
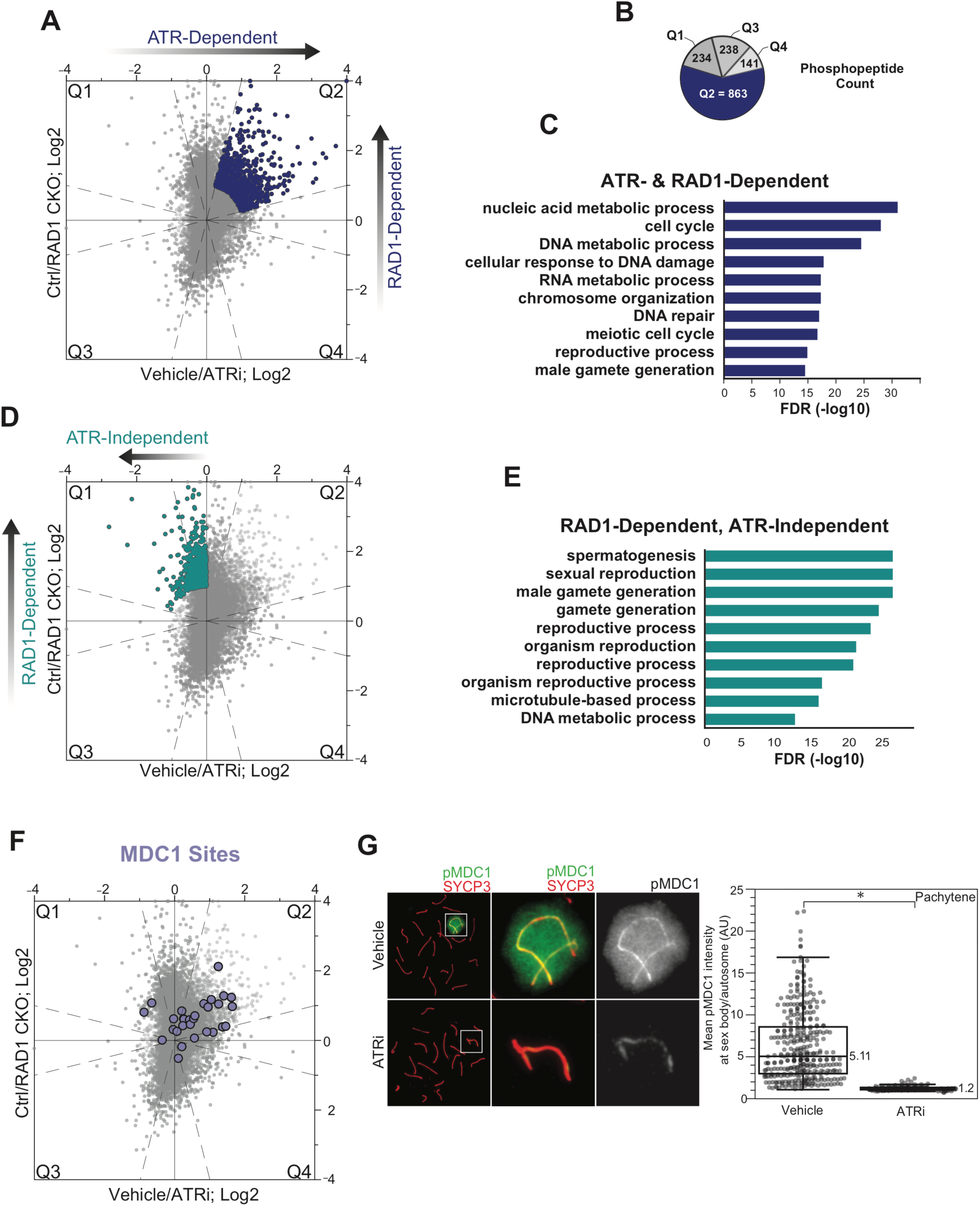
Functional classification analysis of identified signaling events. A) Scatter plot indicating designated set of ATR and RAD1-dependent phosphopeptides (quadrant 2; Q2). Dashed lines indicate filtering thresholds applied to Q2. B) Phosphopeptide count per quadrant. C) Curated gene ontology list of proteins with ATR and RAD1-dependent phosphorylation identified in Q2. D) RAD1-dependent and ATR-independent phosphopeptides in Q1 (no thresholds applied). E) Gene ontology of proteins with RAD1-dependent, ATR-independent phosphorylation. F) Scatter plot highlighting the detected MDC1 phosphopeptides. G) Immunofluorescence of meiotic spreads from mice treated with vehicle or 50 mg/kg AZ20 for four hours and stained for pMDC1 (green) and SYCP3 (red) with quantification of pachytene-staged cells (4 vehicle mice, n=360 cells; 4 ATRi mice, n=282 cells p=0.0268 measured by student’s t-test) (see methods for details)

Depletion of RAD1 in spermatocytes results in reduction in tubule size, infertility and loss of germ cells^42^. Meiotic spreads from *Rad1* CKO mice further showed mid-pachytene arrest^42^. Therefore, we performed gene enrichment analysis on the subset of RAD1-dependent, ATR-independent sites which are depicted in Q1 (Fig. 2D). Our analysis revealed enrichment in functional groups related to spermatogenesis, gamete generation and reproductive processes (Fig. 2E, S4B, table S2). This finding is consistent with our rationale that Q1 would contain phosphorylation events that reflect indirect or pleiotropic events caused by long-term depletion of RAD1 and loss of 9-1-1-mediated ATR signaling. Furthermore, these pleiotropic events are expected to reflect the impairment of processes downstream of meiosis I. Gene ontology of RAD1-independent, ATR-dependent phosphosites found in Q3 (Fig S4A-B, table S2) revealed enrichment for many GO terms involved in cytoskeletal organization and protein polymerization/depolymerization. These targets are potentially regulated by RAD1-independent ETAA1 activation of ATR or reflect events in non-meiotic cell types.

To further assess the quality of the generated dataset, we examined the distribution of phosphopeptides of MDC1, a known ATR target during prophase I that binds to phosphorylated H2AX and further stimulate the spreading of ATR and DNA repair factors on chromatin loops of the X and Y chromosomes to promote MSCI^43, 49^. Thirty MDC1 phosphorylation sites were detected, including ten sites in Q2 (ATR- and RAD1-dependent) and twenty sites that were not dependent on ATR phosphorylation (Fig. 2F-G,S4F). These data reveal that MDC1 is a multiply-phosphorylated, with both ATR-dependent and independent modes of regulation. Furthermore, the identification of MDC1 phosphosites whose phosphorylation status did not change upon ATR inhibition suggested that phosphorylation changes in Q2 are not due to changes in MDC1 protein abundance, nor are they due to non-specific changes in MDC1 phosphorylation status. Notably, we were able to detect at least one phosphorylation site that did not change between the control and reduced ATR activity condition (non-regulated) in over 60% of the proteins containing a phosphorylation site detected in Q2 (Table S1), which indicates that the majority of phosphorylation in Q2 are not likely due to protein abundance changes, but rather regulated and specific phosphorylation events. Nonetheless, for proteins whose phosphorylation sites are only detected in Q2, it remains possible that protein abundance change may be the underlying cause of the observed change in phosphopeptide abundance. Overall, these results validate our experimental rationale and reinforce the importance of combining data from ATRi and *Rad1* CKO datasets to map primary meiosis-specific ATR signaling.

### Connectivity analysis defines ATR-regulated sub-networks

To examine the network of ATR signaling in our dataset, we performed a Cytoscape/ClueGO analysis to systematically define a comprehensive map of processes involving ATR- & RAD1-dependent phosphorylation. ClueGO analysis revealed several expected categories including nucleic acid metabolic processes, regulation of cell cycle, chromosome organization and DNA repair. Additionally, STRING analysis of all proteins with ATR- & RAD1-dependent phosphorylation revealed a densely connected sub-network of DNA repair proteins (Fig. 3A). The sub-network of DNA repair proteins defined by string analysis comprises several proteins involved in ATR recruitment and activation, including TOPBP1, ATRIP, and RAD9B. This sub-network also included a number of key proteins involved in homologous recombination and related DNA repair activities/transactions, such as RAD50, NBS1 (Nbn), CTIP (RBBP8), RAD51C, PALB2, RAD18, SLX4, RAP80 (UIMC1) and RNF168 (Fig. S4F-G). Notably, STRING and ClueGO analysis also identified RNA metabolic processes such as mRNA processing, transcription and splicing as represented groups for proteins identified in Q2 (Fig. 3A, S4C-D). In particular, a sub-network of proteins involved in splicing, including several components of the pre- and post-splicing multi-protein mRNP complexes (Fig. S4C-D) were found. Surprisingly, meiotic ATR signaling also impinged on a highly connected sub-network of proteins involved in centrosome assembly and function, several of which are categorized under the biological process “cell cycle” (Fig. S4E). Overall, these data reveal the scope of processes affected by meiotic ATR signaling, which while extensive, seems to preferentially converge on the control of DNA repair, mRNA processing and cell cycle processes.

**Figure 3.**
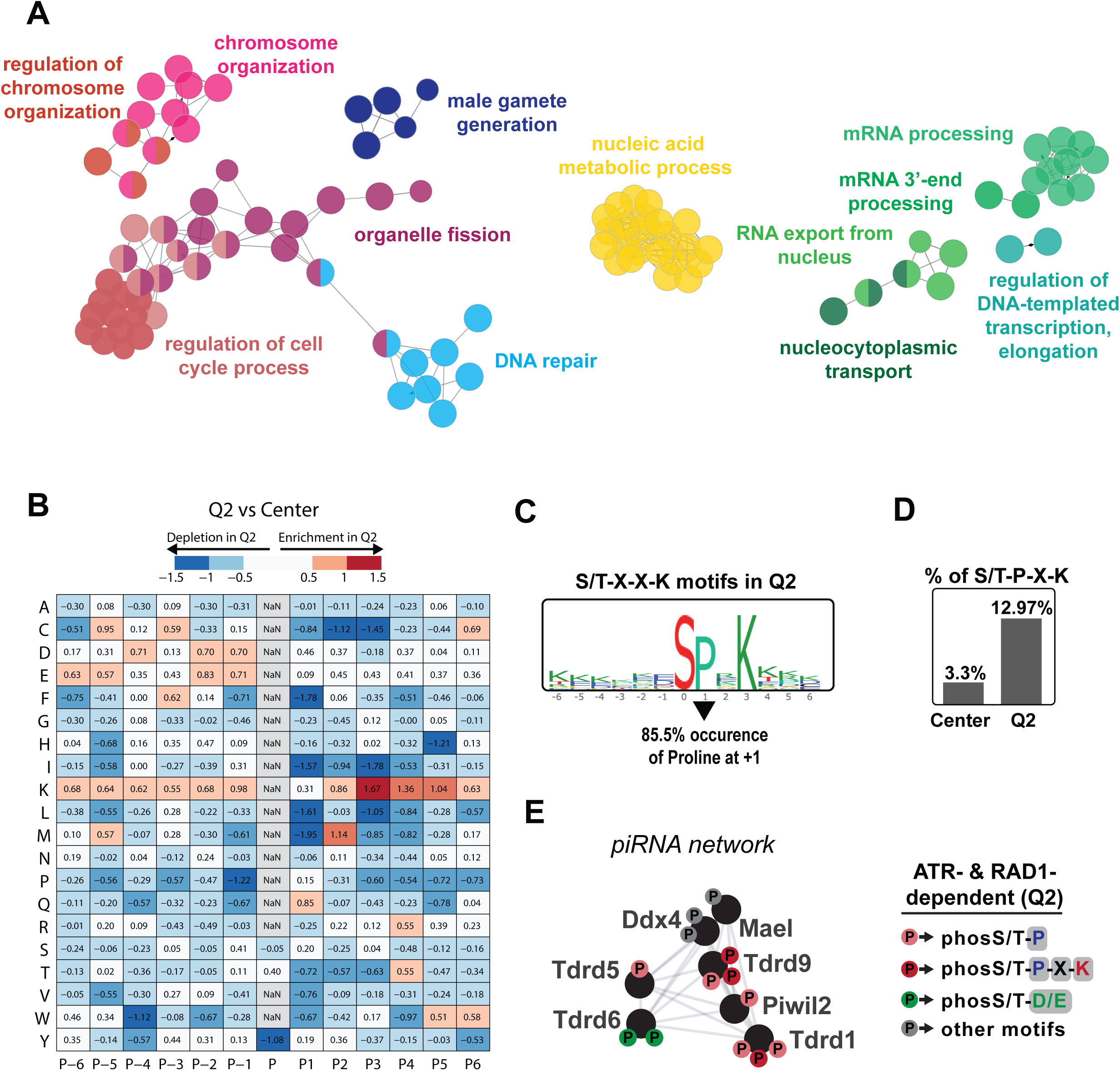
ClueGO and phospho-motif analysis of ATR and RAD1-dependent events in Q2. A) Functional GO network generated by ClueGO analysis of RAD1 and ATR-dependent phosphopeptides in Q2. GO functional groups are separated by color and colored text where nodes with multiple colors belong to multiple GO functional groups. Each node represents a GO term with a P value of <0.05 with size of the node corresponding to the significance of the enrichment. B) Unbiased analysis of amino acid prevalence surrounding identified phosphorylation sites. Prevalence data for Q2 phosphopeptides were compared to prevalence data for all non-regulated phosphopeptides (“Center” of scatter plot in Figure 2) to yield relative depletion or enrichment indexes. Negative values indicate a specific amino acid is less prevalent in phosphopeptides in Q2, and positive values indicate a specific amino acid is more prevalent in phosphopeptides in Q2. “P” indicates position with respect to identified phosphorylation site. RAD1- and ATR-dependent signaling (Q2) is enriched in phosphorylation sites within an S/T-X-X-K motif. C) Analysis of amino acid prevalence in Q2 phosphopeptides having the S/T-X-X-K motif. Larger letters indicate higher prevalence. Image indicates results for serine phosphorylated phosphopeptides. Similar result is obtained for threonine phosphorylated phosphopeptides. Overall occurrence of proline at +1 position for both serine and threonine phosphorylated phosphopeptides in Q2 at S/T-X-X-K motif is 85.5%. D) Bar graph of percentage of S/T-P-X-K motif in the Center (non-regulated) and Q2 regions. E) Schematic of the network of piRNA proteins having ATR- and RAD1-dependent phosphorylation events (Q2). Color represents distinct phospho-motifs.

### Meiotic ATR promotes extensive phospho-signaling at an S/T-P-X-K motif

To further examine the network of RAD1 and ATR-dependent phosphorylation events, we performed an unbiased analysis of phosphorylation motifs in Q2. As shown in Fig. 3B, we computed the relative proportion of each amino acid at the +/-6 positions surrounding the identified phosphorylation sites, comparing their prevalence in Q2 (ATR and RAD1-dependent sites) versus center (unregulated or not-differentially phosphorylated sites). The resulting matrix revealed the relative degree of depletion or enrichment for each amino acid in each position. Notably, the preferred ATR motif (Q at +1 position) was only slightly enriched in Q2. Of the 863 ATR and RAD1 dependent sites identified in Q2, 42 sites were at the S/T-Q consensus motif, a ∼5% prevalence that represents a ∼2-fold enrichment over the prevalence of this motif in the group of unregulated sites (Table S1). The finding that most phosphorylation sites in Q2 are not in the S/T-Q motif suggests that ATR is able to phosphorylate other motifs and/or directly or indirectly regulate the activity of other kinases or phosphatases during meiosis. Consistent with the latter hypothesis, several kinases were found to contain an ATR and RAD1-dependent phosphorylation site, including CDK1/2, MAK, NEK1 and PKMYT1. Strikingly, in addition to the expected enrichment of Q at the +1 position we noticed a drastic enrichment of K at the +3 position. Close inspection of the group of ATR and RAD1-dependent phospho-sites with K at +3 revealed that 85.5% of them contained a P at the +1 position (Fig. 3C). We next compared the prevalence of S/T-P-X-K motifs Q2 sites (ATR- & RAD1-dependent) to the set of unregulated phosphosites found in the center of the dataset and found Q2 had ∼4-fold more S/T-P-X-K. (Fig. 3D, table S1). These results suggest that ATR is activating one or more kinases that have a preference for S/T-P-X-K motif. The identity of the kinase(s) responsible for phosphorylating the large set of ATR- & RAD1-dependent sites (112 sites in Q2) at the S/T-P-X-K remains unknown.

Interestingly, we noticed that several components of the piRNA network were enriched for RAD1- and ATR-dependent phosphorylation within the S/T-P-X-K motif (Fig. 3E). piRNAs are abundant in spermatocytes and protect genome integrity by preventing retrotransposon integration during meiosis^50, 51^. Male mice deficient for piRNA biogenesis such as the those deficient for PIWI proteins are infertile due to spermatocyte arrest^52^. Despite the importance of piRNAs in spermatogenesis, a thorough mechanistic understanding of piRNA regulation and function remains unknown. Several components of the piRNA biogenesis pathway contained one or more non-S/T-Q phosphorylation site. For example, we identified two phosphorylated S/T-P-X-K motifs as well as one S/T-P motif in TDRD9 (Fig. 3E), an RNA helicase that functions in piRNA metabolism and is important for spermatogenesis^53^, suggesting that TDRD9 may be regulated by kinases in a RAD1-and ATR-dependent manner. Although it is unclear which kinase(s) are phosphorylating TDRD9 or other components of the piRNA network, these data implicate the ATR signaling cascade in regulating the piRNA pathway. Further work will be important to dissect the role of ATR in regulating piRNA proteins.

### RAD1- and ATR-dependent phosphorylation at S/T-Q sites defines potentially direct ATR targets involved in DNA damage signaling, DNA repair and RNA metabolism

Given that ATR preferentially phosphorylates S/T-Q motifs^54, 55^, we reasoned that most phosphorylation sites at S/T-Q in Q2 are more likely to reflect direct ATR substrates in meiosis. As expected, proteins involved in DNA damage signaling and repair were found to contain Q2 S/T-Q phosphorylation, including ATR itself, TOPBP1, RAP80, and components of the MRN-CTIP complex (Fig. 4D-E). The group of S/T-Q sites in Q2 also included proteins involved in RNA metabolism and chromatin regulation (Fig. 4B, D-E). Notably, S/T-Q sites in proteins involved in RNA metabolism (SETX, XPO5 and RANBP3) displayed the highest ATR-dependency (Fig. 4E). Most proteins with an S/T-Q phosphorylation site identified in Q2 have additional phosphorylation sites that did not change with RAD1 CKO or ATR inhibition (Fig. 4A-B), indicating that the overall abundance of these proteins was not likely changing. Furthermore, many Q2 proteins contained both an S/T-Q motif and another non-S/T-Q phosphorylation site that was also ATR-dependent (Fig. 4A-B). Taken together, these results reveal a set of proteins that may potentially be direct substrates of ATR in meiosis and define a set of RNA regulatory proteins subjected to 9-1-1 and ATR-dependent phosphoregulation.

**Figure 4.**
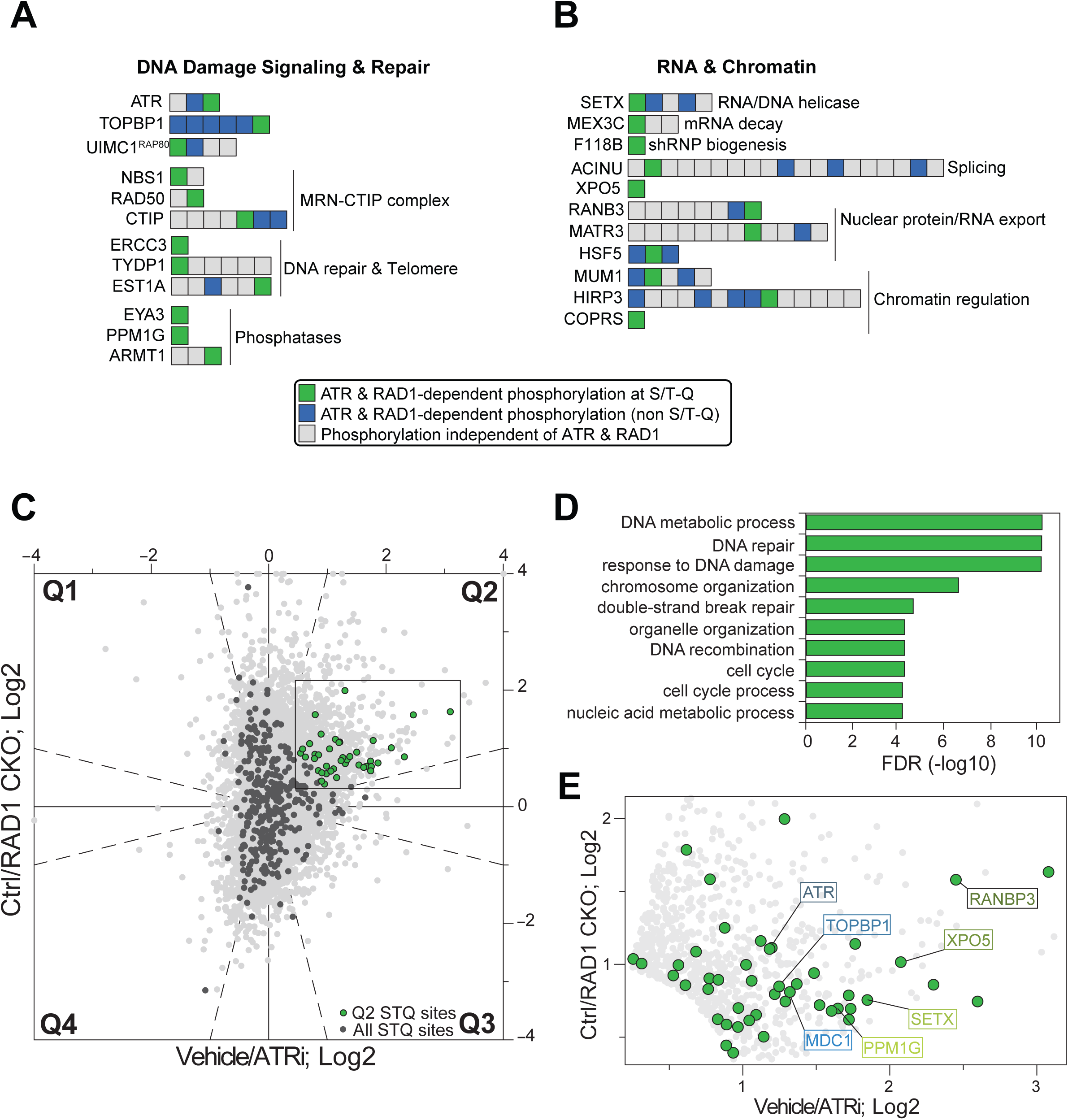
RAD1 and ATR-dependent phosphorylation at the S/T-Q motif includes proteins involved in DNA and RNA processes. A) Selected set of proteins involved in DNA damage signaling and DNA repair found to have RAD1- and ATR-dependent phosphorylation at the S/T-Q motif. Identified phosphorylation sites are ordered sequentially from the n-terminus to the c-terminus of each protein. B) Selected set of proteins involved in chromatin modification and RNA metabolic processes found to have RAD1- and ATR-dependent phosphorylation at the S/T-Q motif. Identified phosphorylation sites are ordered sequentially from the n-terminus to the c-terminus of each protein. C) Scatter plot highlighting all S/T-Q phosphorylation outside Q2 (dark grey) and S/T-Q phosphorylation inside Q2 (green). D) Gene ontology of proteins containing S/T-Q phosphorylation within Q2. E) A zoomed region of the scatter plot in 4C highlighting Q2 S/T-Q phosphopeptides in proteins involved in DNA repair and RNA metabolism.

### ATR modulates the localization of RNA regulatory factors Senataxin and RANBP3

Although ATR localizes to the sex body to promote MSCI, it is not known if ATR directly regulates RNA metabolic proteins to promote silencing or processing of RNAs. To investigate how meiotic ATR may regulate RNA metabolism we focused on RNA metabolic proteins with S/T-Q phosphosites in Q2. We found serine 353 in Senataxin (SETX), an RNA:DNA helicase with established roles in transcriptional regulation and genome maintenance^56^, to be downregulated upon RAD1 loss and ATR inhibition (Fig. 4B and 4E). Senataxin disruption is associated with male infertility in humans and *Setx^-/-^* male mice are infertile resulting from arrest at the pachytene stage in meiosis I^57, 58^. Senataxin localizes to the XY chromosomes and promotes the localization of ATR, yH2AX and other DNA repair and checkpoint factors to the sex body to promote MSCI^59^. Senataxin interacts with many proteins involved in transcription and is thought to regulate multiple aspects of RNA metabolism such as splicing efficiency and transcription termination in part by its activity in resolving R-loops (RNA-DNA hybrids)^60^. To assess whether ATR modulates Senataxin function in meiosis, we stained for Senataxin in meiotic spreads derived from both ATR inhibitor treated and *RAD1*-CKO mice. In accordance with previous work, we found that Senataxin localizes to the sex body at pachynema control spreads^58, 59^. Strikingly, Senataxin accumulation at the sex body was significantly reduced in pachytene spreads derived from ATRi treated mice (Fig. 5A-B, S2A-B). While it is difficult to morphologically distinguish the X and Y chromosomes in the *Rad* CKO spreads, we observed no enrichment of Senataxin around any selection of chromosomes in pachytene-like spreads with four or more synapsed autosomes (Fig.5 C-D,S2C-D). Previous studies have found that Senataxin inhibition results in diminished ATR signaling at the sex body, implicating Senataxin in promoting ATR signaling^58, 59^. Our results further suggest that ATR-dependent phosphorylation promotes the recruitment or retention of Senataxin at the sex body, consistent with a model in which Senataxin and ATR act in a feed-forward loop to cooperatively promote their recruitment and efficient sex body formation and MSCI.

**Figure 5.**
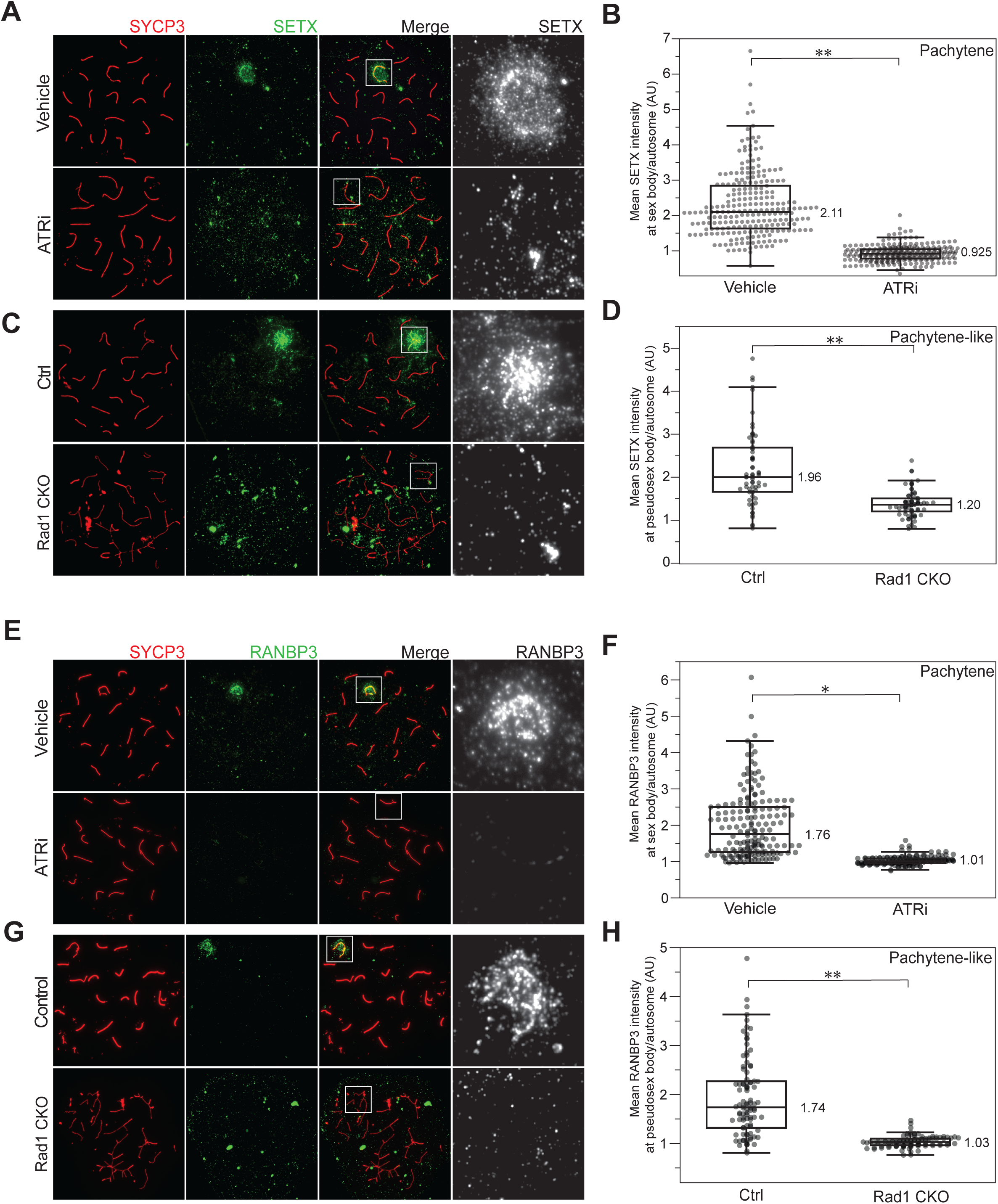
SETX and RANBP3 localization to the sex body is lost upon ATR inhibition. A) Immunofluorescence of meiotic chromosome spreads with SETX (green) and SYCP3 (red) from mice collected 4 hours after 50 mg/kg treatment with AZ20 or vehicle. B) Quantification of pachytene spreads from 5A (4 vehicle mice; n=237 cells; 4 ATRi mice; n=283 cells p=0.00435 measured by student’s t-test). C) Immunofluorescence of meiotic chromosome spreads with SETX (green) and SYCP3 (red) from *Rad1* CKO and control spreads. D) Quantification of pachytene or pachytene-like spreads from 5C (4 control mice, n=64 cells; 4 RAD1 CKO mice, n=72 cells p=0.00286 measured by student’s t-test). E) Immunofluorescence of meiotic chromosome spreads with RANBP3 (green) and SYCP3 (red) from mice collected 4 hours after 50 mg/kg treatment with AZ20 or vehicle. F) Quantification of pachytene spreads from 5E (3 vehicle mice, n=174 cells; 3 ATRi mice, n=167 cells p=0.048 measured by student’s t-test). G) Immunofluorescence of meiotic chromosome spreads with RANBP3 (green) and SYCP3 (red) from *Rad1* CKO and control spreads. H) Quantification of pachytene or pachytene-like spreads from 5G (4 control mice, n=96 cells; 4 CKO mice, n=99 cells p=0.0039 measured by student’s t-test).

Another protein with a S/T-Q phosphorylation site identified in Q2 was RANBP3 (serine 283). RANBP3 is a relatively unknown protein with connections to miRNA and protein export in mitotic cells^61^ . Unfortunately, it is not known if RANBP3 depletion results in a loss of fertility although one study has found an association with decreased RANBP3 expression and human infertility^62^. We investigated the localization of RANBP3 in meiotic spreads and found that in cells derived from wild type or vehicle treated mice, RANBP3 localizes to the sex body at pachynema (Fig 5E-F, S3A-B). The accumulation of RANBP3 is significantly lost at the sex body derived from ATRi treated mice and at all chromosome cores in *Rad1* CKO pachytene-like spreads (Fig 5G-H, S3C-D), suggesting a role for ATR in the recruitment or retention of RANBP3 at the sex body. Overall, these results support a model whereby ATR promotes the proper localization of SETX and RANBP3 to the sex body in pachynema.

## Discussion

ATR has well established roles in promoting genome stability in mitotic cells by regulating multiple aspects of DNA metabolism such as DNA repair, DNA replication and the DNA damage checkpoint^37, 63^. Several phosphoproteomic databases have been generated to characterize the targets of ATR during conditions of replication stress or within the context of mitosis^32, 63–69^. These resources have been useful not only to mechanistically dissect the different roles of ATR, but also to gain a more comprehensive understanding of its multifaceted action in genome metabolism. In the context of meiosis, much less is understood about ATR signaling, and although previous reports have catalogued phosphorylation events in mouse testis using phosphoproteomics, these datasets lack experimentally established kinase-substrate relationships^70–74^. Given the utmost importance of defining the ATR-mediated signaling events in mammalian meiosis to mechanistically dissect its function and mode of action, here we performed an in-depth phosphoproteomic analysis of ATR signaling in meiosis. The success of our work mostly relied on a two-part approach for identifying high-confidence ATR-dependent phosphorylation events. By combining the datasets from the *Rad1* CKO genetic mouse model and ATR inhibitor treated mice, we enhanced our confidence in identifying meiosis-specific ATR functions. As validation of our dataset, we detected known ATR targets such as MDC1 and TOPBP1 in the set of ATR and RAD1-dependent signaling events. Additionally, we observed the expected enrichment for DNA metabolism, DNA repair and cell cycle gene ontology categories. We anticipate that this database will be a useful resource for the meiosis community for further study into the mechanisms of meiotic ATR activity.

The set of ATR-mediated signaling events detected included phosphorylation in proteins involved in RNA metabolism, such as Senataxin and RANBP3. It is tempting to speculate that these are direct targets since they were phosphorylated at the S/T-Q motif and their localization to the XY body was compromised during ATR inhibition. We propose that these data highlight a need for ATR in regulating distinct aspects of RNA metabolism such as R-loop accumulation, splicing, termination and RNA export at the X and Y chromosome to allow for proper silencing. The model implicates ATR as a central regulator of multiple aspects of RNA metabolism during meiosis, and it is interesting that several proteins involved in the piRNA network were also found to be regulated in an ATR-dependent manner. The connection of ATR to RNA metabolism is not completely surprising since other reports on the mitotic action of ATR have uncovered similar connections^75^. Notably, given that silencing of the XY is inextricably linked to prophase I progression, it is likely that the connection of meiotic ATR signaling to RNA metabolism is even more relevant compared to its mitotic signaling. An interesting model to be explored in future work is that SETX and RANBP3 may both play a key coordinated role in removing RNA from XY DNA, and exporting it, to establish MSCI (Fig. S3F). Further work will be needed to establish these as direct ATR substrates and to dissect the mechanism by which ATR promotes the localization and action of Senataxin, RANBP3 and other RNA metabolic proteins identified in this study.

Since our phosphoproteomic is unbiased and not only directed at the preferred S/T-Q motif, we were able to capture a range of phosphorylation events in other motifs suggesting that ATR regulates multiple downstream kinases during meiosis. Strikingly, we observed a strong enrichment for ATR-dependent phosphorylation sites at the S/T-P-X-K motif, suggesting that ATR promotes activation of one of multiple downstream proline-directed kinases. We identified several kinases subjected to ATR and RAD1-dependent phosphorylation, including CDK1/2, MAK, NEK1 and PKMYT1. Of those, CDK1/2 and MAK have established preferential phosphorylation motif consistent with the S/T-P-X-K motif. A simple model would predict that these kinases are activated by ATR during prophase I, which could be tested by future phosphoproteomic analysis of testes from mice treated with inhibitors for these kinases. It is worth mentioning that in mitosis, the canonical action of ATR in promoting DNA damage checkpoint, and consequent cell cycle arrest, is mediated via inhibition CDK activity, and consequent reduction in S/T-P phosphorylation sites^76^. In this sense, the observed dependency of S/T-P-X-K motif for ATR in meiosis is the opposite to what would be predicted from mitotic cells. Since the high activity of ATR in meiosis does not result in meiotic arrest, but is actually required for meiotic progression, it is possible that our data is revealing a drastic difference in how ATR signaling is wired with downstream kinases in meiotic versus mitotic cells. In addition to S/T-P-X-K motif, several other motifs were represented in the set of ATR- and RAD1-dependent sites. We cannot exclude that the presence of other motifs in Q2 is due to the function of ATR regulation of phosphatases. Importantly, several phosphatases, including PPM1G and PP1R7, were also identified in Q2.

Overall, our work represents an initial attempt to reveal the scope of targets and processes affected by meiotic ATR signaling. As expected, the ATR signaling network in meiosis is overwhelming complex and multifaceted. Major challenges will be to untangle the functional relevance of most of the identified signaling events, and understand how the different modes of ATR signaling are coordinated for proper control of meiotic progression. Of importance, ATR may be activated in a 9-1-1-independent manner, via ETAA1. Therefore, additional phosphoproteomic analyses from mouse mutants/CKOs of *ETAA1* are likely to identify different subsets of ATR targets that may represent different modes of ATR signaling in meiosis. Another key outstanding question is to understand how the ATR kinase, which imposes cell cycle checkpoints in most other cell types, is so highly activated in spermatocytes without inducing cell cycle arrest. A potential explanation may lay at the specificity of ATR’s action at the sex body, which may be devoted to the regulation of checkpoint-independent processes such as the control of RNA processing during meiotic prophase I, as supported by our data. Finally, there are medical implications of understanding ATR signaling in meiosis, since many ATR inhibitors are currently in phase 2 clinical trials for cancer treatment and determining the impact of these inhibitors in meiotic cells will be relevant to define the effects of these treatments in patient fertility.

## Materials and Methods

### ATR inhibitor treatment of mice

AZ20 was reconstituted in 10% DMSO (Sigma), 40% propylene glycol (Sigma), and 50% water. Control mice were treated with 10% DMSO (Sigma), 40% propylene glycol (Sigma), and 50% water. Wild-type C57BL/6 male mice aged to 8 weeks-old were gavaged with 50mg/kg of AZ20 (Selleckchem) and euthanized at indicated time points. Specific timepoints examined in this study include collection after 3 days of 50mg/kg, 2.5 days of 50mg/kg per day or 4 hours after one dose of 50mg/kg AZ20. All mice used for this study were handled following federal and institutional guidelines under a protocol approved by the Institutional Anima Care and Use Committee (IACUC) at Cornell University.

### Testes phosphopeptide enrichment and TMT labeling

Whole, decapsulated testes were collected and frozen at -80° C from 8 week-old AZ20 and vehicle-treated C57BL/6 mice 4 hours after treatment as indicated. Whole, decapsulated testes from *Rad1* CKO and littermate control mice were collected at 8 or 12 weeks of age. *Rad1* CKO control genotypes are Rad1^-/Fl^;Cre^-^, Rad1^+/Fl^; Cre^-^ and Rad1^+/Fl^;Cre^+^. Individual testes were thawed at 4° C in lysis buffer (50mM Tris pH 8.0, 5mM EDTA, 150mM NaCl, 0.2% Tergitol) supplemented with 1mM PMSF and PhosSTOP (sigma) and sonicated. 4 mg of protein (quantified by Bradford protein assay, Biorad) was collected, denatured with 1% SDS and reduced with 5mM DTT at 65° C for 10 minutes followed by alkylation with 60mM Iodoacetamide. Proteins were precipitated in a cold solution of 50% acetone, 49.9% ethanol and 0.1% acetic acid and protein pellet was washed once with a solution of 0.08M Urea in water. Protein pellet was resuspended at a concentration of 10mg/ml in 8M urea, Tris 0.05M, pH 8.0, NaCl 0,15M and diluted 5-fold prior digestion with 80ug of trypsin (TPCK-treated, Sigma) overnight at 37° C. Protein digests were acidified to a final concentration of 1% TFA and cleaned-up in a solid-phase extraction (SPE) C_18_ cartridge pre-conditioned with 0.1% TFA solution. Peptides were eluted from the cartidges with 80% acetonitrile, 0.1% acetic acid aqueous solution and dried in speed-vac. Phosphopeptide enrichment was performed using a Thermo-Fisher Fe-NTA phosphopeptides enrichment kit according to the manufacturer protocol (Cat# A32992, ThermoScientific). Phosphopeptides samples were split into 4 aliquots (10%, 30%, 30% and 30%) and dried in silanized glass tubes. For each experiment, the three 30% aliquots from each control and AZ20-treated or control and RAD1-KO samples were resuspended in 35uL of 50mM HEPES and labeled with 100ug of each of the TMTsixplex Isobaric Label Reagents (ThermoFisher), previously diluted in 15uL of pure acetonitrile. TMT-labeling reaction was carried out at room temperature for 1h and quenched with 50uL of 1M Glycine. All 6-plex TMT labeled aliquots were mixed, diluted with 200uL of aqueous solution of formic acid 1% (v/v) and cleaned up in a SPE 1cc C_18_ cartridge (Sep-Pak C18 cc vac cartridge, 50 mg Sorbent, WATERS). Bound TMT-labeled phosphopeptides were eluted with 50% acetonitrile, 0.1% formic acid in water and dried in a speed-vac.

### Mass spectrometric analysis of TMT-labeled phosphopeptides

Dried TMT-labeled phosphopeptides were ressuspended in 16.5uL water, 10uL formic acid 10% (v.v) and 60uL of pure acetonitrile and submitted to HILIC pre-fractionation. Samples were fractionated using a TSK gel Amide-80 column (2 mm x 150 mm, 5 μm; Tosoh Bioscience), using a three-solvent system: buffer A (90% acetonitrile), buffer B (75% acetonitrile and 0.005% trifluoroacetic acid), and buffer C (0.025% trifluoroacetic acid). The chromatographic runs were carried out at 150uL/min and gradient used was: 100% buffer A at time = 0 min; 94% buffer B and 6% buffer C at t = 3 min; 65.6% buffer B and 34.4% buffer C at t = 30 min with a curve factor of 7; 5% buffer B and 95% buffer C at t = 32 min; isocratic hold until t = 37 min; 100% buffer A at t = 39 to 51 min. One-minute fractions were collected between minutes 8 and 10 of the gradient; 30-second fractions between minutes 10 and 26; and two-minute fractions between minutes 26 and 38 for a total of 40 fractions. Individual fractions were combined according chromatographic features, dried in speedvac and individually submitted to LC-MS/MS analysis. Individual phosphopeptide fractions were resuspended in 0.1% trifluoroacetic acid and subjected to LC-MS/MS analysis in an UltiMate™ 3000 RSLC nano chromatographic system coupled to a Q-Exactive HF mass spectrometer (Thermo Fisher Scientific). The chromatographic separation was carried out in 35-cm-long 100-µm inner diameter column packed in-house with 3 µm C_18_ reversed-phase resin (Reprosil Pur C18AQ 3μm). Q-Exactive HF was operated in data-dependent mode with survey scans acquired in the Orbitrap mass analyzer over the range of 380 to 1800 m/z with a mass resolution of 120,000. MS/MS spectra was performed selecting the top 15 most abundant +2, +3 or +4 ions and precursor isolation window of 0.8 m/z. Selected ions were fragmented by Higher-energy Collisional Dissociation (HCD) with normalized collision energies of 38 and the mass spectra acquired in the Orbitrap mass analyzer with a monitored first mass of 100m/z, mass resolution of 60,000, AGC target set to 1×10^5^ and max injection time set to 110ms. A dynamic exclusion window was set for 30 seconds.

### Phosphoproteomic data analysis

The peptide identification and quantification pipeline relied on MaxQuant platform (v.1.6.3.4) and the Andromeda search engine^77^. The Mouse UNIPROT proteome database (22,297 entries) was downloaded on 2018-10-22 and supplemented with the default MaxQuant contaminant protein database. Search parameters included tryptic requirement, 20 ppm for the precursor match tolerance, dynamic mass modification of 79.966331Da for phosphorylation of serine, threonine and tyrosine and static mass modification of 57.021465Da for alkylated cysteine residues. TMT-labeling correction parameters were added according the information provided by manufacturer. All the additional MaxQuant parameters were the default. The original raw files representing each set of fractions for each experiment were organized into groups for data processing. Phospho(STY)Sites.txt MaxQuant output table was processed in order to obtain the TMT ratio for the 7 individual experiments, for each phosphosite identified. Additional sets of criteria were applied to select for high confidence phosphosite identification and regulation, as presented in results sections. The phosphoproteomic data generated in this study were deposited to the Massive database (http://massive.ucsd.edu) and received the ID: MSV000086764, doi:10.25345/C57N54, and ProteomeExchange ID: PXD023803.

### Meiotic chromosome spread immunofluorescence

Meiotic spreads were produced as previously described^78^. Briefly, decapsulated testes tubules were incubated in a hypotonic extraction buffer (30mM Tris pH 7.2, 50mM sucrose, 17mM citrate, 5mM EDTA, 0.5mM DTT, 0.1mM PMSF) for 1 hour. 1mm sections of tubule were dissected in 100mM sucrose solution and then added to slides coated in 1% paraformaldehyde/0.15% Triton-X and allowed to spread for 2.5 hours in a humidification chamber. Slides were then dried for 30 minutes and washed in 0.4% photoflo (Kodak)/PBS solution for 5 minutes. Slides were immediately processed for immunofluorescence or frozen at -80°C. For staining, slides were washed in a solution of 0.4% photoflo/PBS for 10 minutes, 0.1% Triton X/PBS for 10 minutes and blocked in 10% antibody dilution buffer (3% BSA, 10% goat serum, 0.0125% Triton X)/PBS for 10 minutes. Primary antibodies were diluted in antibody dilution buffer at the indicated dilution and incubated with a strip of parafilm to spread the antibody solution in a humidification chamber at 4°C overnight. After primary antibody incubation, the slides are washed with 0.4% photoflo/PBS for 10 minutes, 0.1% Triton-X for 10 minutes and blocked with 10% antibody dilution buffer. Secondary antibodies were diluted as indicated and incubated on slides with a parafim strip at 37°C for 1 hour. Slides were then washed in 0.4% photoflo/PBS for 10 minutes twice followed by 0.4% photoflo/H_2_O for 10 minutes twice and allowed to dry before mounting with DAPI/antifade.

### Imaging and quantification

Slides were imaged on a Leica DMi8 Microscope with a Leica DFC9000 GTC camera using the LAS X (Leica Application Suite X) software. For every condition, a minimum of 50 images from three independent mice were acquired. To quantify florescence intensity, the LAS X software quantification tool was used. Briefly, a ROI (region of interest) line was drawn over the sex body and mean intensity of the underlying pixels was recorded. Additionally, two ROI lines of equal length were placed over two autosome cores and the mean pixel intensity was also recorded to serve as an internal control for background florescence. The sex body ROI intensity was then normalized to the average of the two autosomal ROI intensities for each individual cell. For the *Rad1* CKO mice, where a true sex body does not form, a line was drawn over chromosomes that were best morphologically identified as the sex chromosomes by SYCP3 staining.

### Antibody list

| Reagent- Primary Antibody | Source | Identifier | Dilution |
| --- | --- | --- | --- |
| yH2AX | Millipore | 05-636 | 1:10000 |
| pMDC1 (phosphoT4) | Abcam | Ab35967 | 1:500 |
| SETX | Abcam | Ab220827 | 1:100 |
| RANBP3 | Bethyl | IHC-00295 | 1:100 |
| SYCP3 (mouse) | Custom<sup>79</sup> | N/A | 1:2000 |
| SYCP3 (rabbit) | Abcam | Ab15093 | 1:2000 |

| Reagent- Secondary Antibody | Identifier | Dilution |
| --- | --- | --- |
| Goat anti-mouse Alexa Fluor 488 | 62-6511 | 1:2000 |
| Goat anti-mouse Alexa Flour 555 | A-10521 | 1:2000 |
| Goat anti-rabbit Alexa Fluor 488 | 65-6111 | 1:2000 |
| Goat anti-rabbit Alexa Fluor 555 | A-10520 | 1:2000 |

## Acknowledgements

We thank all members of the Smolka, Weiss and Cohen Labs for valuable discussions related to this work. The authors thank Robert Gingras and Mason Muir for assistance with imaging and image quantification, Fenghua Hu and Tony Bretscher for use of the microscopes, and Shannon Marshall for assistance with phosphopeptide enrichment. Figure 1A and 2F was created using BioRender.com. This work is supported by a grant from the National Institute of Health (R01-HD095296) to M.B.S. and R.W.

## Author Contributions

MS, RW, JS and CP designed the study. JS and CP performed phosphoproteomic experiments. JS performed all imaging and quantification of meiotic spreads. VF conceived of data processing pipeline. VF and GAM wrote scripts for data analysis. PC and RF provided critical reagents. JS and MS wrote the manuscript. PC and RS edited the manuscript.

---

**Supplemental Figure 1.**
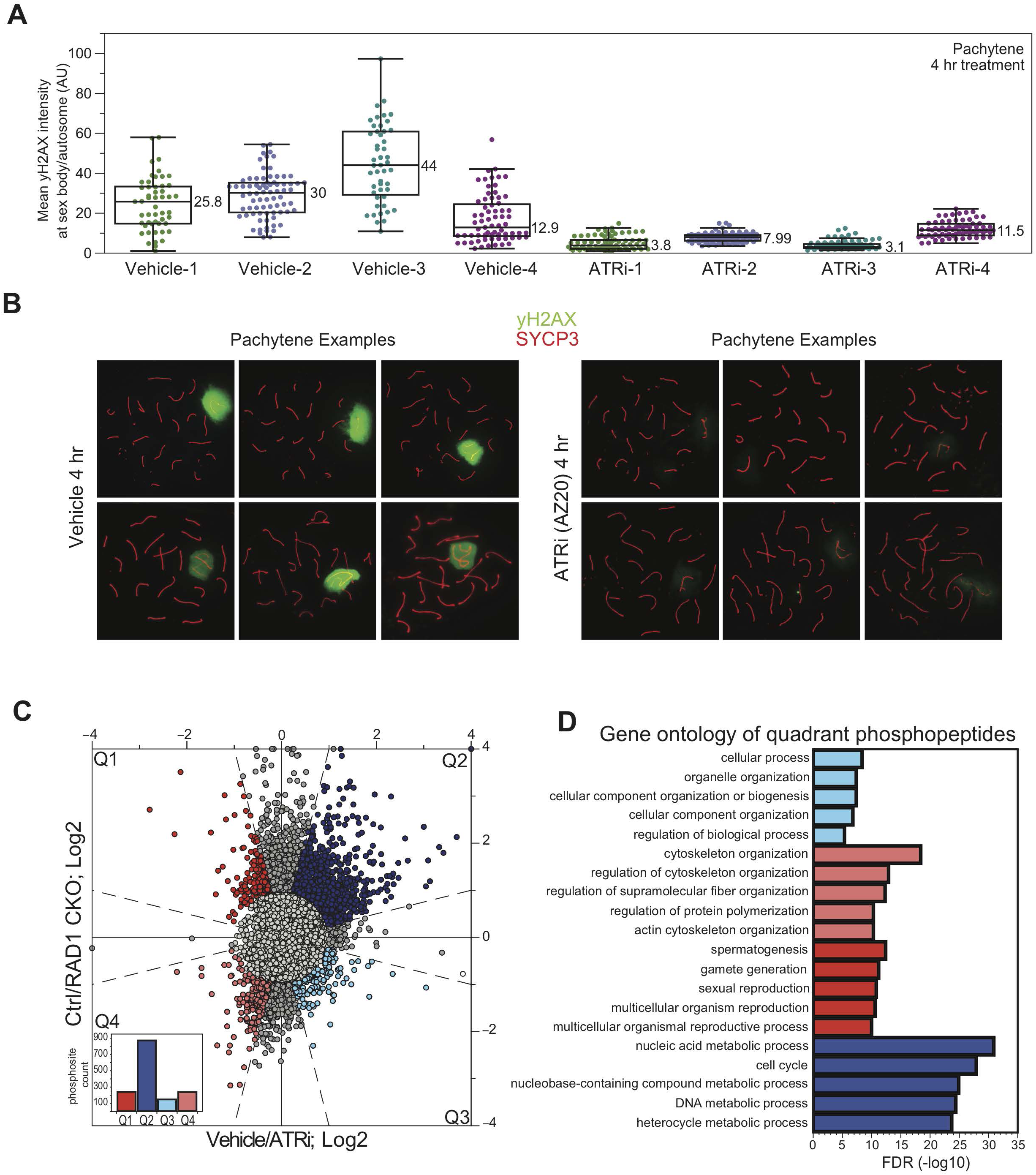
yH2AX is reduced at the sex body after 4 hours of 50 mg/kg treatment with AZ20 and gene ontology of quadrants. a) quantification mean intensity of the ratio of yH2AX signal as depicted in Figure 1C separated by individual animal replicates. yH2AX intensity is measured as described in methods. Datapoints indicate the ratio of signal intensity across sex body to average of intensity across two autosomes for an individual pachytene-stage meiotic spread. b) example of pachytene spreads showing variation in signal intensity and pattern from quantification in S1A with yH2AX (green) and SYCP3 (red). c) scatterplot identifying peptides in each quadrant with phosphopeptide count (bar graph, lower left) and corresponding top five STRING analysis categories in d) bar graph.

**Supplemental Figure 2.**
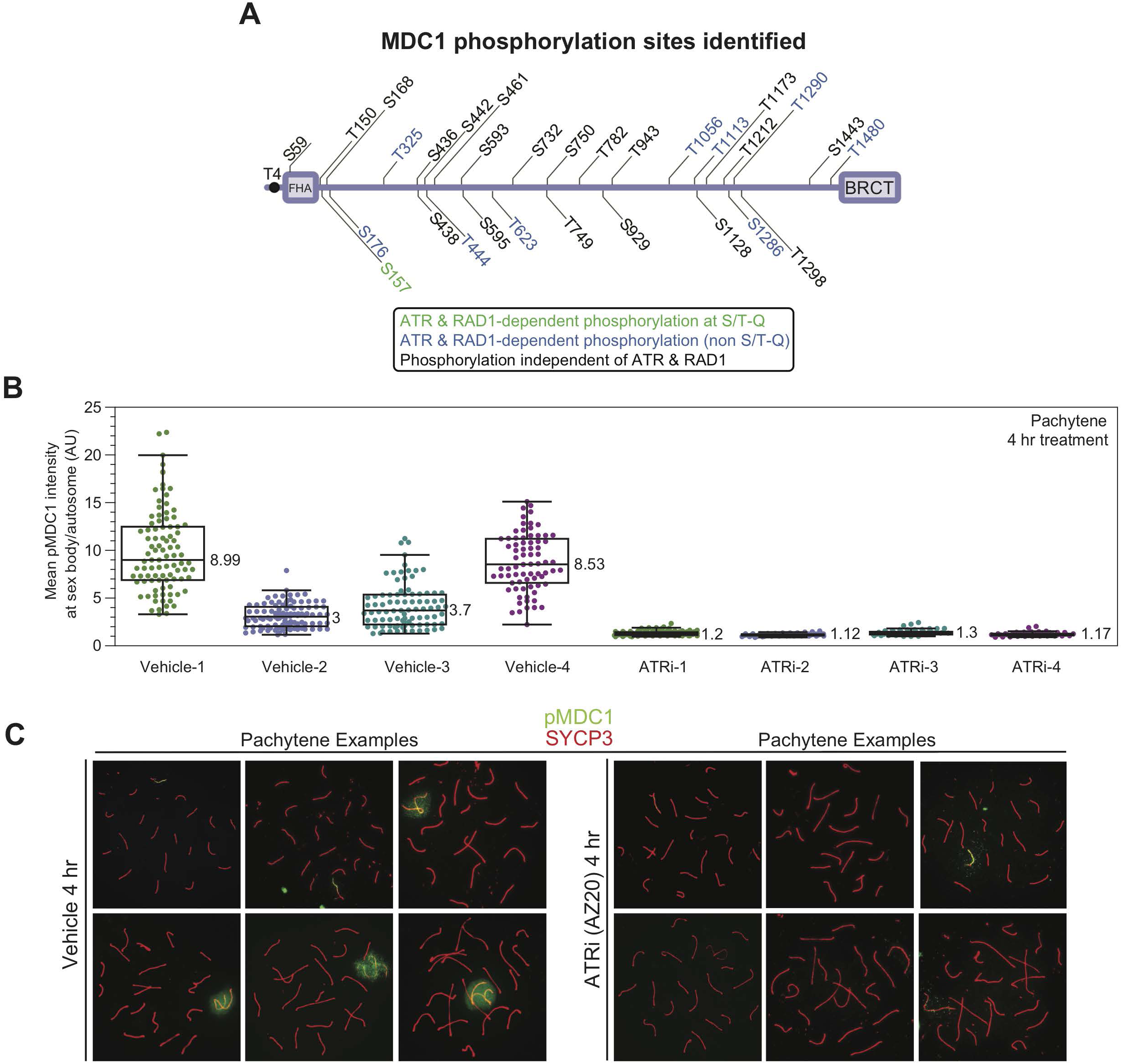
pMDC1 signal is lost at the sex body upon ATR inhibition. a) Schematic of detected phosphorylation sites in MDC1 b) quantified pMDC1 intensity from 2G separated by individual mice after treatment with vehicle or ATR inhibitor for 4 hours as indicated. Quantification was done as described in methods. c) example meiotic spreads depicting variation in signal intensity and pattern for pMDC1 (green) and SYCP3 (red) for ATRi and vehicle-treated mice.

**Supplemental Figure 3.**
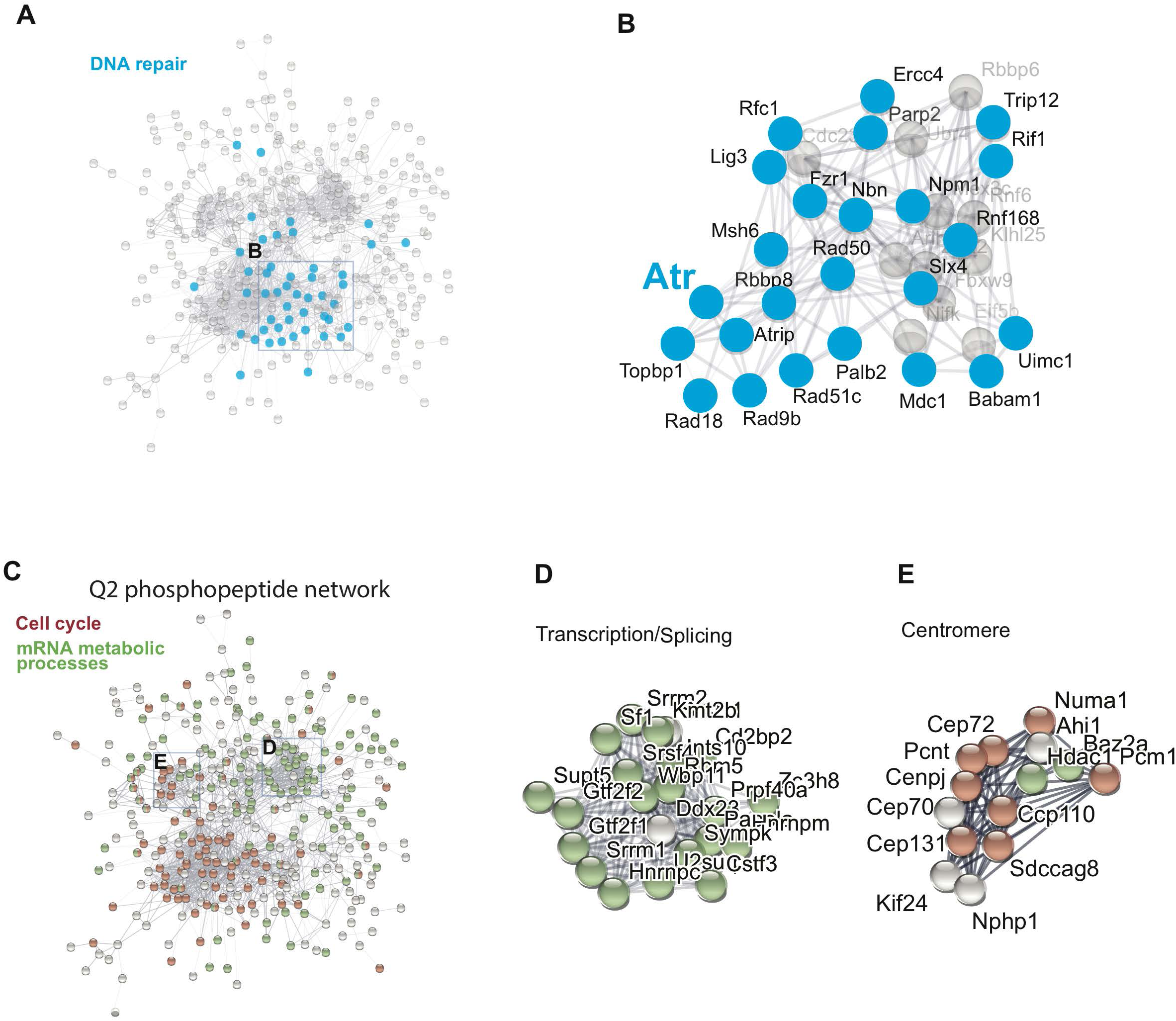
Gene ontology of protein networks. a) STRING analysis of Q2 proteins with network of highlighted proteins in the GO DNA repair category b) detail of DNA repair protein network highlighting ATR-associated node. c) STRING network of cell cycle and mRNA metabolic proteins. d) detail of transcription/splicing and e) centromere proteins

**Supplemental Figure 4.**
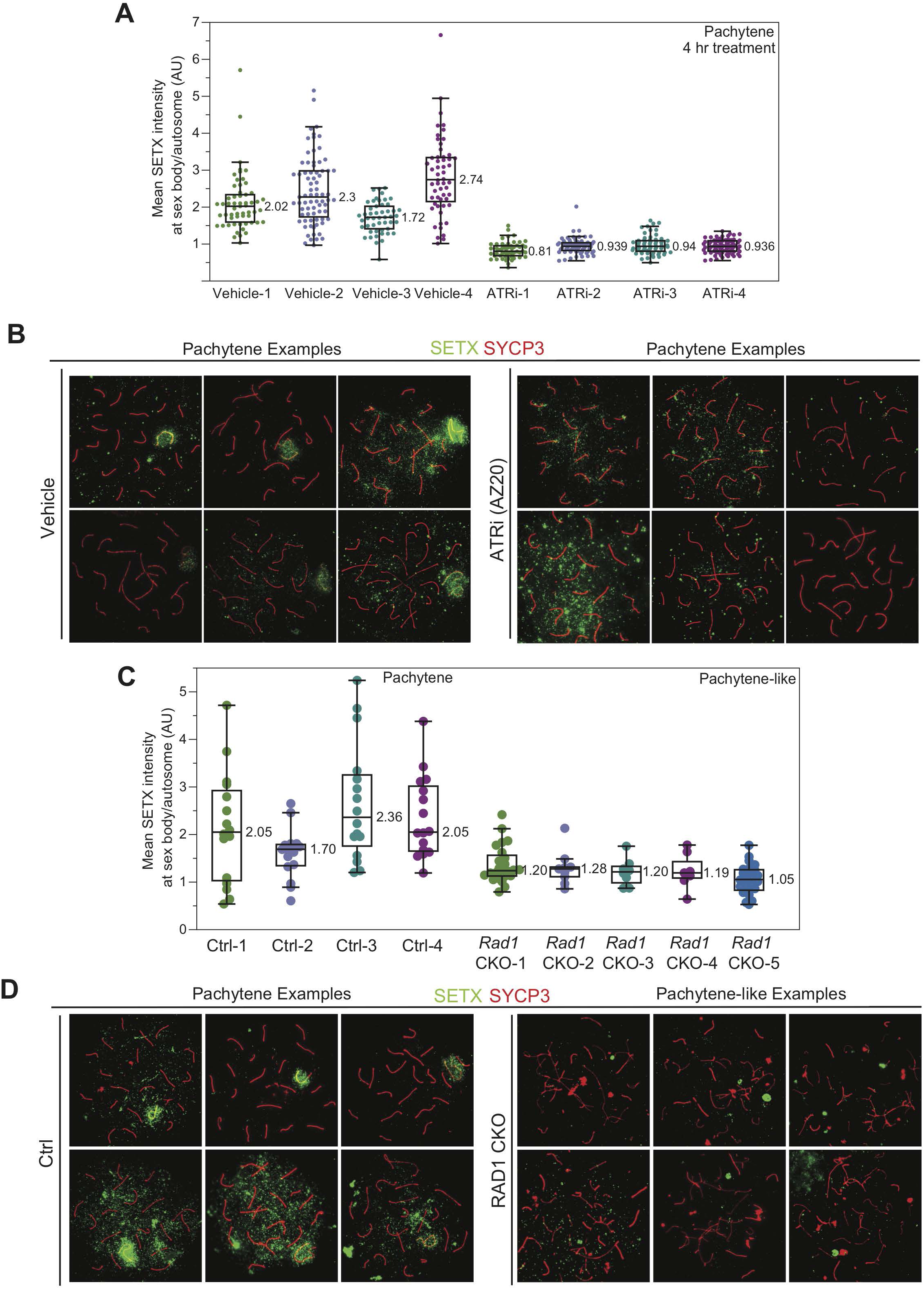
SETX signal is lost at the sex body upon ATR inhibition. a) quantification of SETX signal at the sex body separated by individual mice with c) example images from vehicle or ATR inhibitor treated mice collected 4 hours after 50 mg/kg treatment with AZ20 or vehicle. c) quantification of SETX at sex body or sex chromosomes of control or *Rad1* CKO mice, respectively. d) example spreads of SETX (green) and SYCP3 (red) staining. *Rad1* CKO ‘pachytene-like’ stage was defined as having 3 or more fully synapsed autosomes. See methods for more details on quantification.

**Supplemental Figure 5.**
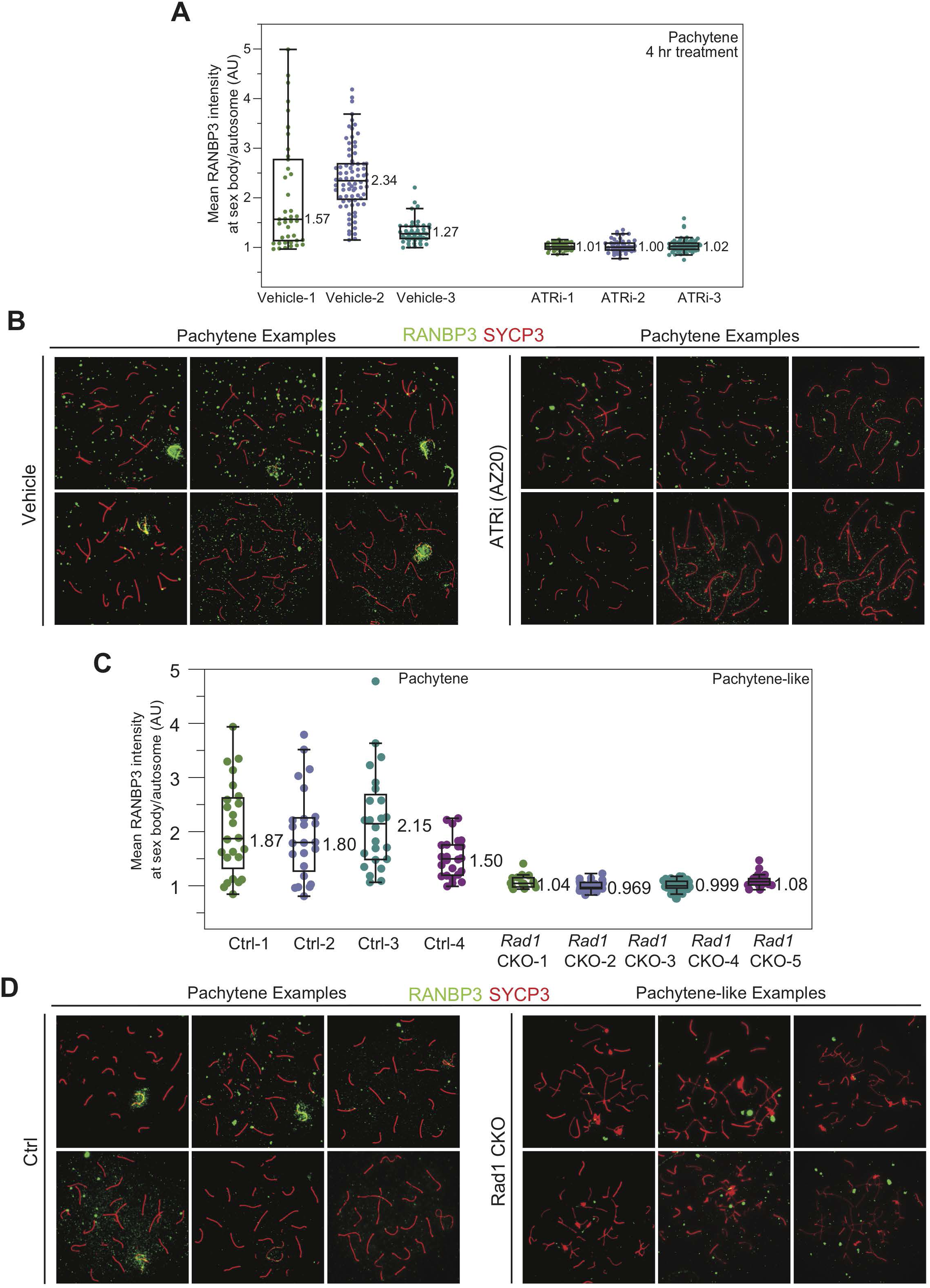
RANBP3 signal is lost at the sex body upon ATR inhibition. a) Quantification RANBP3 intensity from meiotic spreads separated by animal with b) example spreads from mice collected 4 hours after 50 mg/kg treatment with AZ20 or vehicle. c) quantification of RANBP3 at the sex body or sex chromosomes body of control or *Rad1* CKO meiotic spreads, respectively with d) examples images. See methods for more details on quantification.

**Supplemental Figure 6.**
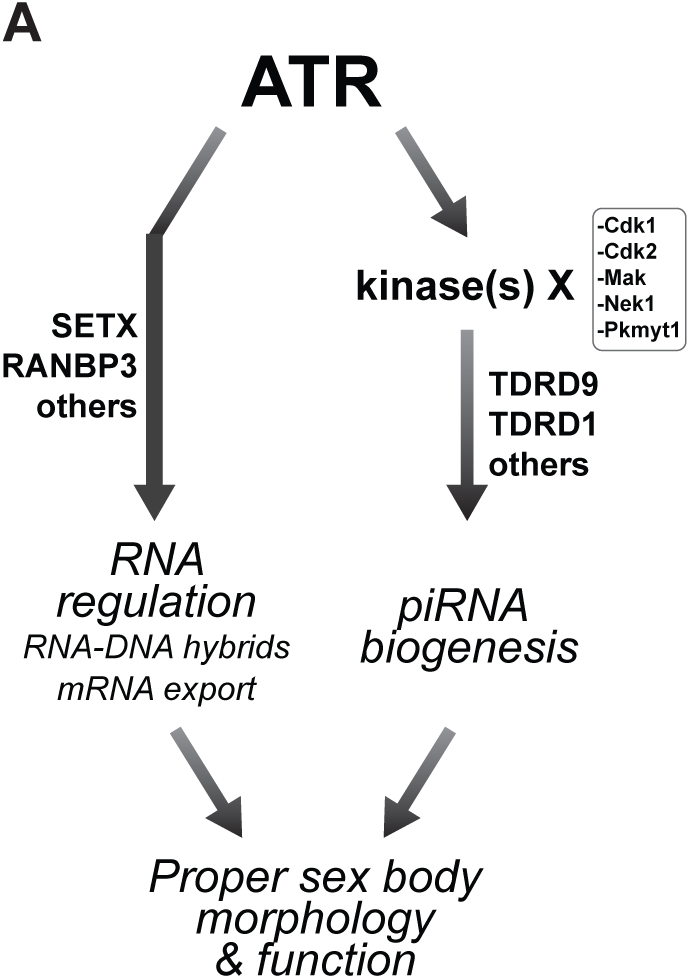
Speculative model for ATR-mediated control of RNA processes during prophase I of mammalian meiosis. Based on the model, ATR directly phosphorylates proteins involved in RNA metabolism, such as SETX and RANBP3, to promote their robust localization to the sex body. In this scenario, sex body-localized SETX and RANBP3 would contribute to ATR-mediated silencing by favoring the disengagement of mRNA from XY chromatin and mRNA export from the sex body. ATR also regulates the activity of additional kinases that in turn phosphorylate piRNA biogenesis-related proteins that may also contribute to sex body morphology and function. Collectively, ATR signaling promotes a range of events contributing to sex body formation.

